# Sugar-mediated inhibition of growth and lignocellulose degradation in anaerobic gut fungi revealed using cellulose filter paper

**DOI:** 10.64898/2026.08.19.745825

**Authors:** Jessica L. Matthews, Stephen C. Fry, Jolanda M. van Munster

## Abstract

Anaerobic gut fungi (AGF) are central to the degradation of plant material in the digestive systems of herbivores. However, how their environment influences their colonisation and degradation of complex biomass is unclear. Here, cellulose filter paper was used as a simplified model of the plant cell wall to investigate how the presence of free sugars in the rumen can affect AGF growth and degradative responses of phylogenetically distinct AGF isolates. From this, galactose was revealed to be inhibitory to both Neocallimastix frontalis and Caecomyces communis, and mannose inhibitory to C. communis. Complete inhibition of C. communis growth was conserved when galactose and mannose were added in their polymeric forms, whereas in contrast, N. frontalis growth was unaffected. This indicates, depending on the AGF isolate, the presence of free sugars and their polymeric form may influence AGF growth through regulatory and metabolic interactions - even if the sugar cannot be utilised for growth as the sole substrate. Collectively, this work highlights the functional diversity in AGF carbohydrate responses and the need for greater understanding of their metabolic regulation for applications in lignocellulosic bioconversion and ruminant nutrition.

## Introduction

Lignocellulose, the structural component of plant cell walls, is a complex structure composed of polymers cellulose, hemicelluloses, pectin, and lignin. As the most abundant raw renewable material on Earth, it is an attractive feedstock for the sustainable production of fuels and chemicals traditionally derived from fossil fuels (Jatoi *et al.,* 2023). However, its recalcitrant nature is a major bottleneck for its efficient conversion, requiring either high energy chemical pre-treatment or relatively slow degradation by a consortium of microbes (Blasi *et al.,* 2023; Jatoi *et al.,* 2023). An ideal candidate to improve the valorisation of lignocellulose are fungi of the phylum Neocallismastigomycota, commonly known as anaerobic gut fungi (AGF) (Hibbett *et al.,* 2007). These fungi possess an arsenal of abundant and highly diverse carbohydrate degrading enzymes (CAZymes) and are amongst the most effective natural degraders of untreated lignocellulose (Kovács *et al.,* 2025). Nevertheless, currently insight in the metabolism is incomplete, limiting their application in both biorefineries and in strategies for increasing productivity in ruminant farming (Kovács *et al.,* 2025).

In their native rumen environment, AGF are the primary colonisers of lignocellulosic material and are integral to the degradation of fibre (Grenet and Barry, 1988; Huws *et al.,* 2021). Compared to aerobic fungi, AGF encode a broad and functionally redundant repertoire of CAZymes (Gruninger *et al.,* 2018) which are generally considered to be less tightly regulated and, in some cases, the expression of genes encoding these enzymes resembles the constitutive expression observed for some anaerobic bacteria (Couger *et al.,* 2015). However, beyond this, it is unclear how the presence of transiently available free sugars in the rumen environment influences AGF regulation of their degradative machinery.

In well-characterised bacterial and fungal systems, it is known that responses to soluble sugars can be mediated through carbon sensing and regulatory systems, such as carbon catabolite repression (CCR) (Adnan *et al.,* 2017). This regulatory system prioritises the metabolism of the preferred carbon source by inhibiting the expression of genes which facilitate the metabolism of alternative substrates (Adnan *et al.,* 2017). AGF belong to an early-diverging fungal lineage (Hanafy *et al.,* 2022) and are distinct from the Dikarya fungi for which these CCR mechanisms are well investigated, but there are some reports in the literature that they too possess CCR (Mountfort and Asher, 1983; Henske *et al.,* 2018a). Furthermore, it has been shown that in some cases micro-organisms can respond physiologically to the presence of sugars even when they are not able to utilise them as carbon sources for growth (Hayer, Stratford, and Archer, 2013; Schmidt and O’Donnell, 2021). As AGF can liberate sugars from complex polysaccharides which they cannot utilise for growth, it is unknown how they influence AGF metabolism and growth responses.

AGF isolates *Neocallimastix frontalis* CoB3 and *Caecomyces communis* SHB have previously been characterised to have distinct degradative effects on lignocellulosic feed composition; *N. frontalis* CoB3 degraded cellulose and hemicellulose to a similar extent, whereas *C. communis* SHB degradation preference depended on the substrate (Shen *et al.,* 2026). Furthermore, both isolates showed differences in preferential uptake of sugars, and *N. frontalis* CoB3 exhibits a co-substrate utilisation of mannose which is dependent on the concentration of glucose (Matthews *et al.,* 2026). Despite these differences, both of their primary metabolisms were identified as being glucose and fructose driven, as they could only utilise a small number of free sugars available in the rumen environment as the sole carbon source (Matthews *et al.,* 2026). This suggests that both *N. frontalis* CoB3 and *C. communis* SHB are able to use cellulose as growth substrate, and that while doing so, they may respond differently to the presence of other carbohydrates.

Here, a simplified model of the plant cell wall was used to investigate how AGF detect and response to simple sugars when growing on ingested plant material in the rumen environment. Filter paper was used as a source for cellulose, the core of lignocellulose, similar what has been used previously to investigate growth and cellulolytic abilities of both AGF (Lowe, Theodorou, and Trinci, 1987; Henske *et al.,* 2018a) and bacteria (Bernalier, Fonty, and Gouet, 1991). To investigate the potential variability in AGF responses, we assayed the phylogenetically distinct isolates *N. frontalis* CoB3 and *C. communis* SHB. This revealed markedly different fungal responses and physical degradation of filter paper, which is almost pure cellulose (Costa *et al.,* 2014; Ferrara *et al,* 2024; Sandy, Manning and Bollet, 2010). However, regardless of these differences, both fungal isolates growth was inhibited by the presence of galactose and *C. communis* SHB was also inhibited by the presence of mannose - despite no measurable responses in growth parameters when glucose was the baseline substrate (Matthews *et al.,* 2026). To further investigate how AGF perceive and respond to sugars in the presence of filter paper, we added monosaccharides and/or polymers of interest to the filter paper. The inhibitory effect of mannose as a free sugar on *C. communis* SHB growth was conserved also when galactomannan and glucomannan were present, but *N. frontalis* CoB3 growth was unperturbed by the addition of these mannan-containing polymers.

Together, these observations indicate that AGF can physiologically respond to the presence of sugars even when they are not able to utilise them as carbon sources for growth. The differing responses of both fungi investigated here highlight the variation of AGF metabolism and degradative capabilities. They also indicate the importance of further investigation to fully elucidated how AGF regulate the synthesis and secretion of their coveted biomass-degrading machineries within the highly competitive microbiome environment of their host digestive system if they are to be effectively implemented in optimising biorefinery and ruminant farming fibre degradation.

## Materials and Methods

### Fungal cultivation

Axenic fungal isolates of *N. frontalis* CoB3 and *C. communis* SHB (Shen *et al.,* 2026) were cultured and maintained under anaerobic conditions in medium C as previously described (Matthews *et al.,* 2026).

Unless stated otherwise, the experimental culture medium was prepared as previously described, with Whatman No. 1 filter paper (1 %, w/v; > 98 %, Sigma-Aldrich) as the carbon source. Where needed, this was supplemented with mono-and disaccharides of lignocellulose (D-glucose, D-xylose, L-arabinose, D-galactose, D-mannose, and D-cellobiose) and those commonly found in the rumen environment (D-maltose and D-lactose), prepared as filter-sterilised stock solutions, to a final concentration of 2.5 g L^-1^. Where required plant-derived polymers (Table 1) were supplemented to 0.5 % (w/v) the weight to volume ratio of additional polymers was chosen through preliminary screening of *N. frontalis* CoB3 growth at varying concentrations of xylan (Fig. S2). Where D-glucose formed the sole carbon source, it was at 5 g L^-1^. All cultures were inoculated with 5 % (v/v) 3-day-old fungal cultures grown on wheat straw (0.5 mm particle size, 1 %, w/v). The production of fermentation gases is used as the proxy for measuring anaerobic fungal growth (Theodorou et al., 1995; Wilken et al., 2020), and culture pH was measured described in (Matthews *et al.,* 2026).

**Table 1:**
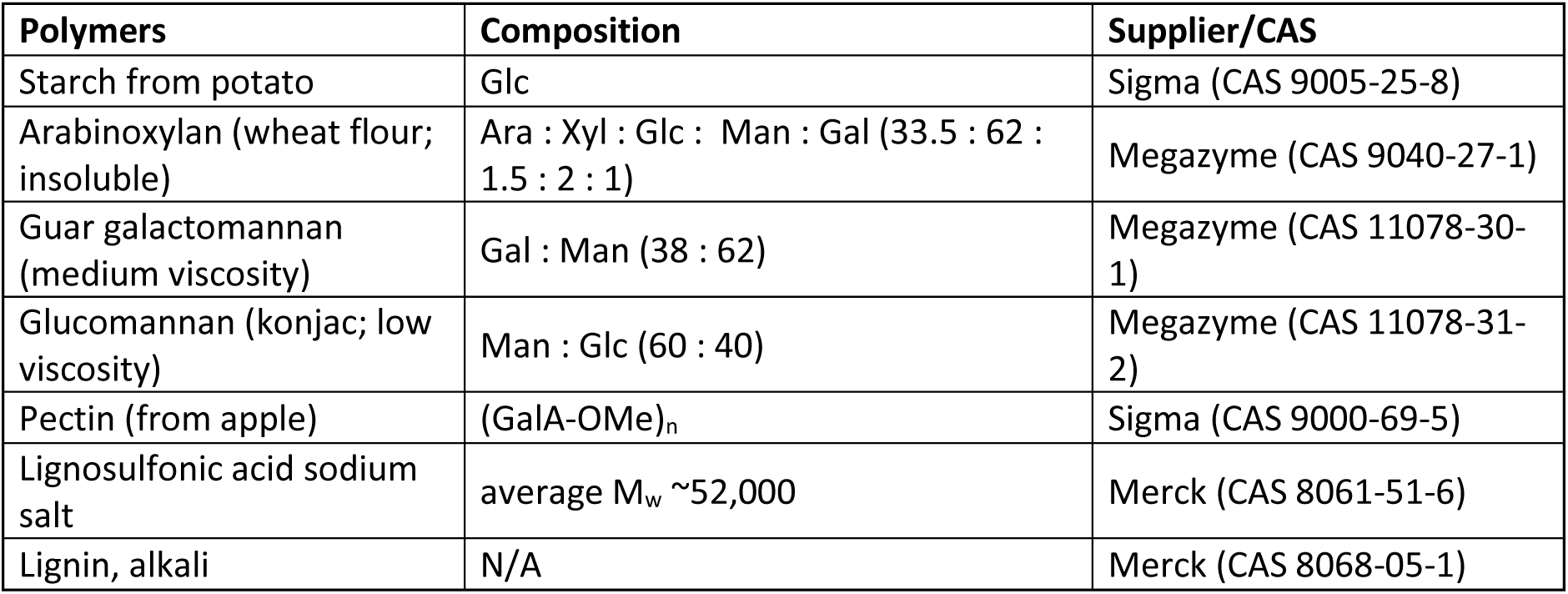
Description of polymers added to filter paper media. Arabinose (Ara), galactose (Gal), glucose (Glc), mannose (Man), xylose (Xyl), galacturonic acid methyl ester (GalA-OMe).

### Thin-layer chromatography

Thin-layer chromatography (TLC) was performed with 3 µL of each culture medium sample and 2.5 µL of standards onto a Merck silica-gel 60 TLC plate. Standards included a cello-oligosaccharide ladder (degree of polymerization, DP 1 to 6, 0.1 %, w/v), or malto-oligosaccharide ladder (DP 1 to 6, 0.1 %, w/v) and 2.5 µL of the additional sugars investigated each at 0.1 % (w/v). The TLC plate was developed in ethyl acetate/pyridine/acetic acid/water (6:3:1:1 by volume), with two ascents. The plate was stained with thymol/H_2_SO_4_/ethanol (0.5:5:95, w/v/v) and heated at 105 °C for 15 minutes (Franková & Fry, 2021).

### Determination of sugar concentration thresholds affecting AGF isolate growth on filter paper

*N. frontalis* CoB3 cultures were supplemented with decreasing concentrations of galactose, whereas *C. communis* SHB cultures were supplemented with decreasing concentrations of galactose or mannose in separate treatments. For each sugar, 50-, 100-, and 1000-fold dilutions of 5 g L^-1^ concentration was used to assess inhibitory thresholds across orders of magnitude. Additional concentrations were selected to also encompass estimated sugar abundances present in commercially available Megazyme polysaccharide preparations, guar galactomannan (medium viscosity) and glucomannan (konjac; low viscosity) (Table 1).

### Addition of inhibitory sugars to established AGF cultures

At the respective exponential growth stage of *N. frontalis* CoB3 and *C. communis* SHB cultures, previously identified (Fig. S1), 0.1 mL of supernatant was removed and immediately placed on ice. Both fungal isolates’ cultures were then subjected to single addition treatments of either galactose (2.5 g L^-1^), mannitol (2.5 g L^-1^ as an osmotic control), or hygromycin (150 µg mL^-1^ as a comparison for growth inhibition). *C. communis* SHB was also treated with mannose (2.5 g L^-1^). Control cultures received no supplementary treatments.

At the end point of the culture, before pH was recorded, 0.1 mL of supernatant was again removed from all cultures and immediately placed on ice. All culture samples were stored at –20 °C until use, when they were thawed at room temperature and centrifuged briefly at 11,000 *g* to pellet any biomass and analysed by TLC.

### Effect of sugar addition on alternative complex biomass carbon sources

*N. frontalis* CoB3 culture media were prepared as described above, with starch (Table 1; 1 %, w/v) and with milled wheat straw (0.5 mm particle size; 1 %, w/v) as the carbon source and supplemented with galactose to a final concentration of 2.5 g L^-1^. *C. communis* SHB culture media were prepared with milled wheat straw (0.5 mm particle size; 1 %, w/v) as the carbon source were supplemented with galactose and mannose respectively, to a final concentration of 2.5 g L^-1^. A 1.0 mL aliquot of each culture was sampled for TLC analysis after the final gas pressure measurement, handled and stored as described above. For both AGF isolates and conditions, three biological replicates displaying the least variation in total gas production were used for TLC analysis.

### Xylosidase enzyme assays

Fungal culture supernatant was obtained by removal of 0.3 mL aliquots from cultures every 24 hours, after the gas pressure had been recorded. Aliquots were briefly centrifuged at 11,000 *g*, pelleting fungal and plant biomass. Samples were assayed immediately for extracellular xylosidase activity, avoiding enzyme denaturation, in a 180-µL reaction mixture containing 2 mM 4-nitrophenyl-β-D-xylopyranoside (xylo-pNP; Megazyme, CAS 2001-96-9), 40 µL culture medium and 80 mM acetate buffer (Na^+^, pH 6.5) in an Eppendorf vial. Each culture medium sample collected was assayed in triplicate. Immediately after the supernatant had been added to the reaction mixture, 40 µL was removed to allow for absorbance correction of the culture medium colour. The remainder of the enzyme reaction mix was incubated at 39 °C for 2 hours in an Eppendorf thermoblock, after which the vials were briefly centrifuged and 40 µL was immediately added to a 96-well plate containing 100 µL sodium carbonate (1 M). The absorbance was measured at 405 nm in the ClarioStar Plus microplate reader, and the concentration of pNP released was calculated using a standard curve of 4-nitrophenol (Sigma, CAS 100-02-7).

### Data Analysis

The rate of exponential growth was estimated from the log_10_ transformed mean accumulated gas pressure of AGF isolates and visual identification of the region of linearity. To this region, linear regression was then applied, and the resulting slope was taken as the exponential growth rate as shown in Figure S1.

Per dataset, each AGF isolate’s respective results were assessed independently for normality using QQ-plot and Shapiro-Wilk. If the normality of residuals correlated linearly and the Shapiro-Wilk test was not significant (*p*-value > 0.05), parametric statistical tests were performed: either Student’s t-test, one-way or two ANOVA (significant *p-*value ≤ 0.05). To assess where the significant differences between means lay if the ANOVA was significant (*p*-value ≤ 0.05), we performed Tukey HSD. If the data were not normally distributed and the Shapiro-Wilk test was significant (*p*-value ≤ 0.05), non-parametric tests were performed: Mann-Whitney U test or the Kruskal-Wallis test followed by pairwise Wilcoxon test if significant (significant *p*-value ≤ 0.05).

## Results

### Fungal isolates have different effects on filter paper

Previous characterisation of soluble sugar utilisation demonstrated that the primary metabolism of both *N. frontalis* CoB3 and *C. communis* SHB is likely to be driven by glucose-and fructose-based substrates (Matthews *et al.,* 2026). Within the AGF isolates’ native environment, the most likely source of glucose is cellulose; therefore, to assess their growth, we used filter paper (which almost pure cellulose) as the primary carbon source. In addition, glucose and wheat straw – a representative lignocellulosic substrate – were used as controls.

For both AGF isolates, the total fermentation gas accumulation when grown on cellulose and wheat straw was comparable, whereas significantly greater gas production was observed when these fungi were grown on glucose (Fig. 1B).

**Figure 1.**
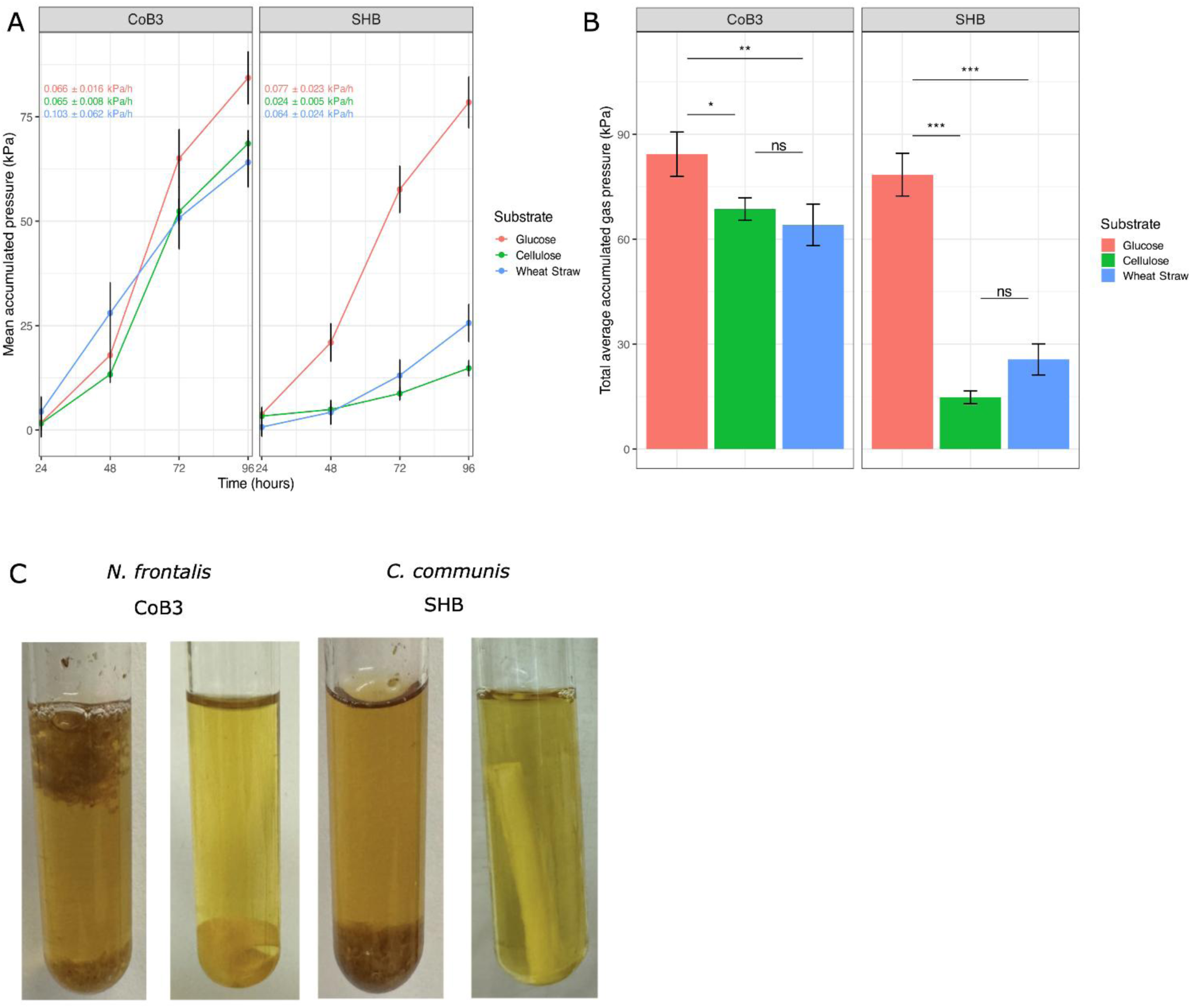
Growth of fungal isolates N. frontalis CoB3 and C. communis SHB on glucose (5 g L^-1^), filter paper and wheat straw. Values reported are mean ± stdev (n=5). A) Fermentation gas accumulation of cultures (graphs) and rate of exponential growth (numerical values in inset; see Fig. S1), B) total fermentation gas accumulated by 96 h. C) A photograph of a representative replicate of each AGF on wheat straw (left) and filter paper (right) at stationary phase (96 h). One-way ANOVA and Tukey HSD analysis was used to assess significance. For all statistical tests p-values are given: *** p < 0.001; ** 0.001 ≤ p < 0.01; * 0.01 ≤ p < 0.05; ns p ≥ 0.05.

*N. frontalis* CoB3 and *C. communis* SHB exhibited markedly different degradation phenotypes on both filter paper and wheat straw. Growth of CoB3 on wheat straw resulted in the formation of a floating biomass raft, where the fungal biomass appeared to aggregate and bind wheat straw particles together. In contrast, SHB growth caused the wheat straw to clump, but to a noticeably lesser extent than observed for CoB3 (Fig. 1D). Most interestingly, there was a vastly different degradation of the filter paper by these isolates, with *N. frontalis* CoB3 curling the filter paper and *C. communis* SHB swelling the filter paper (Fig. 1D).

These results show that filter paper is an ideal model system of the cell wall backbone. Given the markedly different effects observed on the filter paper by these phylogenetically distinct isolates, further investigation was warranted for both *N. frontalis* CoB3 and *C. communis* SHB as they may have different responses in the growth and degradation of cellulose.

### Growth of AGF isolates on filter paper is responsive to the presence of soluble sugars

Previously, we demonstrated that when *N. frontalis* CoB3 and *C. communis* SHB were grown on glucose, additional sugars could repress fungal growth or be taken up and metabolised – even if they could not be used as a sole carbon source (Matthews *et al.,* 2026). Therefore, we performed a similar growth screen to assess the fungal isolates’ responses to the presence of a range of soluble sugars that would be typically available in the natural rumen environment, using filter paper as a primary carbon source. As a source of cellulose, the filter paper provides a more complex and ecologically relevant framework for assessing the effects of soluble sugars on AGF growth in the rumen environment. Furthermore, this screen also allowed us to determine how substrate availability may also influence fungal responses to additional sugars, by comparing glucose (a readily available carbon source) and cellulose (a highly crystalline and insoluble glucose polymer).

Supplementation by soluble sugars elicited vastly different responses from *N. frontalis* CoB3 and *C. communis* SHB. The addition of two sugars known to be readily metabolised by SHB (glucose and cellobiose) significantly increase the total accumulation of fermentation gases and the rate of exponential growth, while the reduced pH of the culture medium indicated an increase in the production of acidic fermentation end products, likely to be VFAs (Fig. 2A,B). In addition, increased physical degradation of the filter paper was observed upon addition of glucose or cellobiose (Fig. 2C). This suggests that these sugars have a beneficial effect on SHB growth; therefore, SHB growth on filter paper alone may be limited by what quantity of metabolisable degradation products it can liberate from the filter paper. Not all sugars known to be metabolisable by SHB had this beneficial effect to SHB growth, as observed the addition of lactose – which is metabolisable by SHB. However, the growth performance of SHB was poor utilising lactose as the sole carbon source (Matthews *et al.,* 2026).

**Figure 2.**
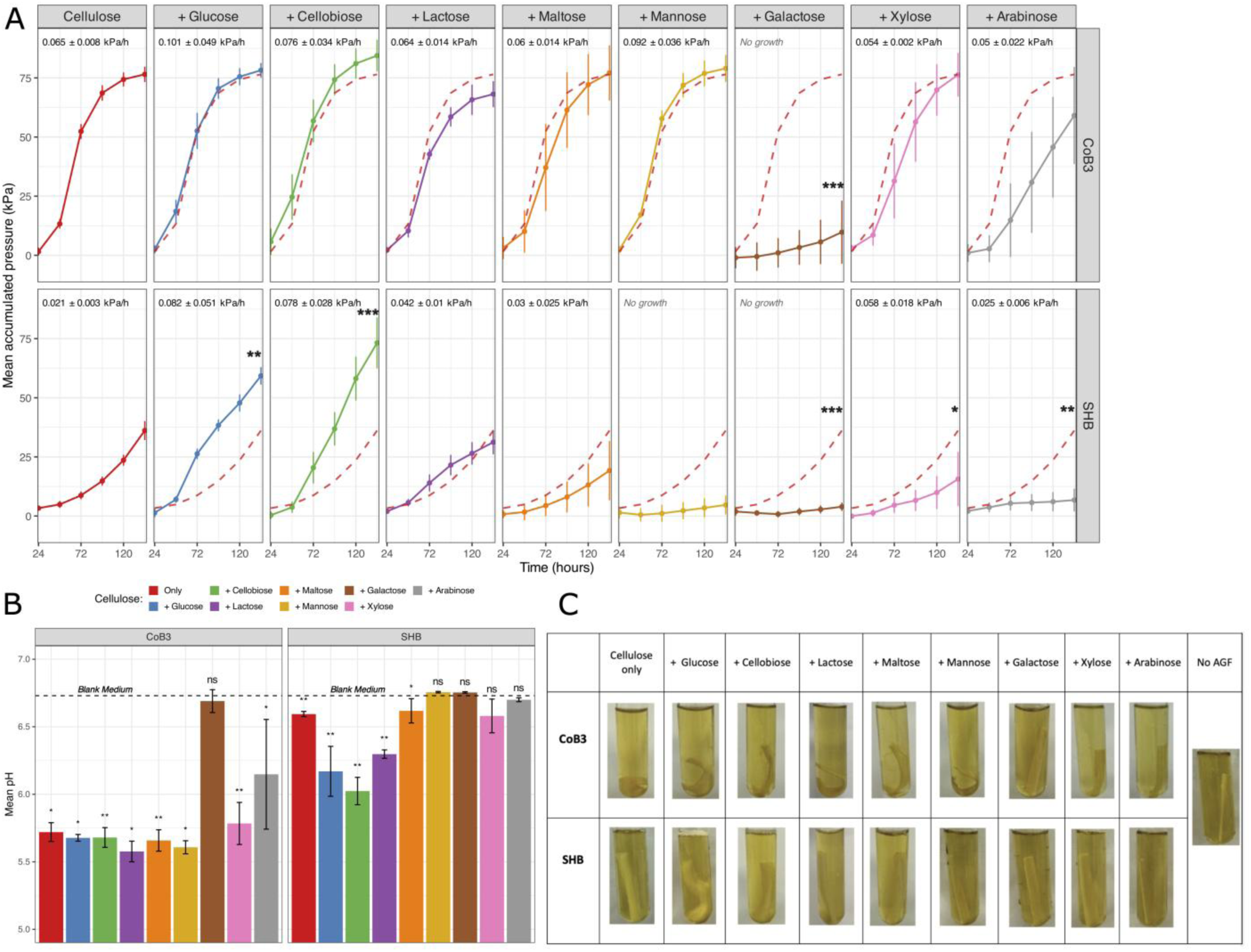
Growth of N. frontalis CoB3 and C. communis SHB on filter paper with additional soluble sugar. Filter paper (1 % (w/v)) supplemented with sugars of interest to a final concentration of 2.5 g L-1. Values reported are mean ± stdev (n=5). A) Fermentation gas accumulation of cultures and rate of exponential growth, B) pH of culture after 144 h after fungal inoculation. C) Photograph of representative culture at 144 h after fungal inoculation. Significance values reported are for AGF isolates on filter paper with additional sugar cultures versus cellulose-only cultures for gas pressure and versus blank medium for pH; one-way ANOVA and Tukey HSD analysis was used to assess both AGF isolates gas pressure and pH of CoB3, and for SHB pH Kruskal-Wallis and pairwise Wilcoxon test was used. For all statistical tests p-values are given: *** p < 0.001; ** 0.001 ≤ p < 0.01; * 0.01 ≤ p < 0.05, ns p ≥ 0.05.

The addition of metabolisable sugars did not significantly affect *N. frontalis* CoB3 growth as measured by fermentation gas accumulation and culture medium pH (Fig. 2A,B). However, the physical degradation of the filter paper seemed to be affected, as the observed curling was less pronounced (Fig. 2C), suggesting a delay in employment of the degradative enzymes to break down the filter paper – indicative of carbon catabolite repression (CCR) (Adnan *et al.,* 2017).

To gain further insight of how the supplementation of metabolisable sugars in the presence of filter paper may have affected AGF growth, the predicted gas pressure of co-substrate conditions was estimated from the sum of measured gas pressure accumulation of the metabolisable sugar (2.5 g L^-1^) and filter paper, when each were the sole carbon source (Fig. S3). This predicted gas pressure was compared to that of the corresponding experimentally measured co-substrate condition, to determine if the presence of the free sugar significantly reduced the experimentally measured fermentation gas production – indicative of CCR. From this, the experimentally measured gas pressure accumulation of *N. frontalis* CoB3 was significantly reduced compared to the respective predicted gas pressure accumulation, as well as that of *C. communis* SHB when supplemented with glucose or lactose respectively (Fig. S3). However, the experimental total gas accumulation of SHB for cellulose and cellobiose was not statistically different compared to the predicted (Fig. S3), suggesting a beneficial effect of cellobiose for growth on cellulose, perhaps priming SHB’s enzymatic machinery for cellulose degradation.

The supplementation of non-metabolisable sugars significantly reduced *C. communis* SHB growth; both the pentose sugars arabinose and xylose significantly reduced the accumulation of fermentation gases, arabinose more so than xylose (Fig. 2A). Unexpectedly, supplementation with the hexose sugars mannose and galactose resulted in complete SHB growth inhibition - no accumulation of fermentation gas was recorded (Fig. 2A) and no changes in pH compared to the blank control medium (Fig. 2B), nor were any physical changes of the filter paper observed (Fig. 2C). This complete inhibition of growth by galactose was also observed for *N. frontalis* CoB3 (Fig. 2), where it was the only non-metabolisable sugar to significantly affect CoB3 fermentation output and to inhibit degradation or curling of the filter paper (Fig. 2).

In general, the growth of *C. communis* SHB isolate was more sensitive to the presence of soluble sugars than that of *N. frontalis* CoB3, which may reflect a greater responsiveness to sugar-mediated signalling within its ecological niche. The parallel observation that galactose supplementation resulted in complete inhibition of *N. frontalis* CoB3 and *C. communis* SHB suggests a potential shared underlying response, perhaps linked to common features in metabolic pathways involved in sensing, transporting, or initiating degradation of more complex polymeric substrates.

### Low concentrations of galactose and mannose affect growth of N. frontalis and C. communis on cellulose

Whilst a high concentration of sugar may reflect locally elevated concentrations present during rapid plant material degradation, typically the sugar concentration in the rumen is considered to be variable and spatially heterogeneous (Nikkhah, 2014). To understand the sensitivity of the inhibitory effects of galactose (on *N. frontalis* CoB3 and *C. communis* SHB) and mannose (on *C. communis* SHB), and to contextualise if this inhibition could be potentially observed in the rumen or anaerobic digestors – we assessed the concentration threshold for growth inhibition.

*C. communis* SHB growth was extremely sensitive to the presence the hexose sugars even at the lowest concentration tested, 0.005 g L^-1^, as shown by the significantly reduced fermentation gas accumulation upon galactose and mannose addition (Fig. 3). Isolate SHB was more sensitive to the presence of mannose, compared to galactose, as the threshold of complete growth inhibition was observed at a lower concentration (0.156 g L^-1^ and 0.313 g L^-1^ for mannose and galactose respectively; Fig. 5). At all concentrations screened, there was an observed absence of filter paper swelling (Fig. 1D) which previously was observed to correlated with successful *C. communis* SHB growth using filter paper as the primary carbon source (Fig. 1).

**Figure 3.**
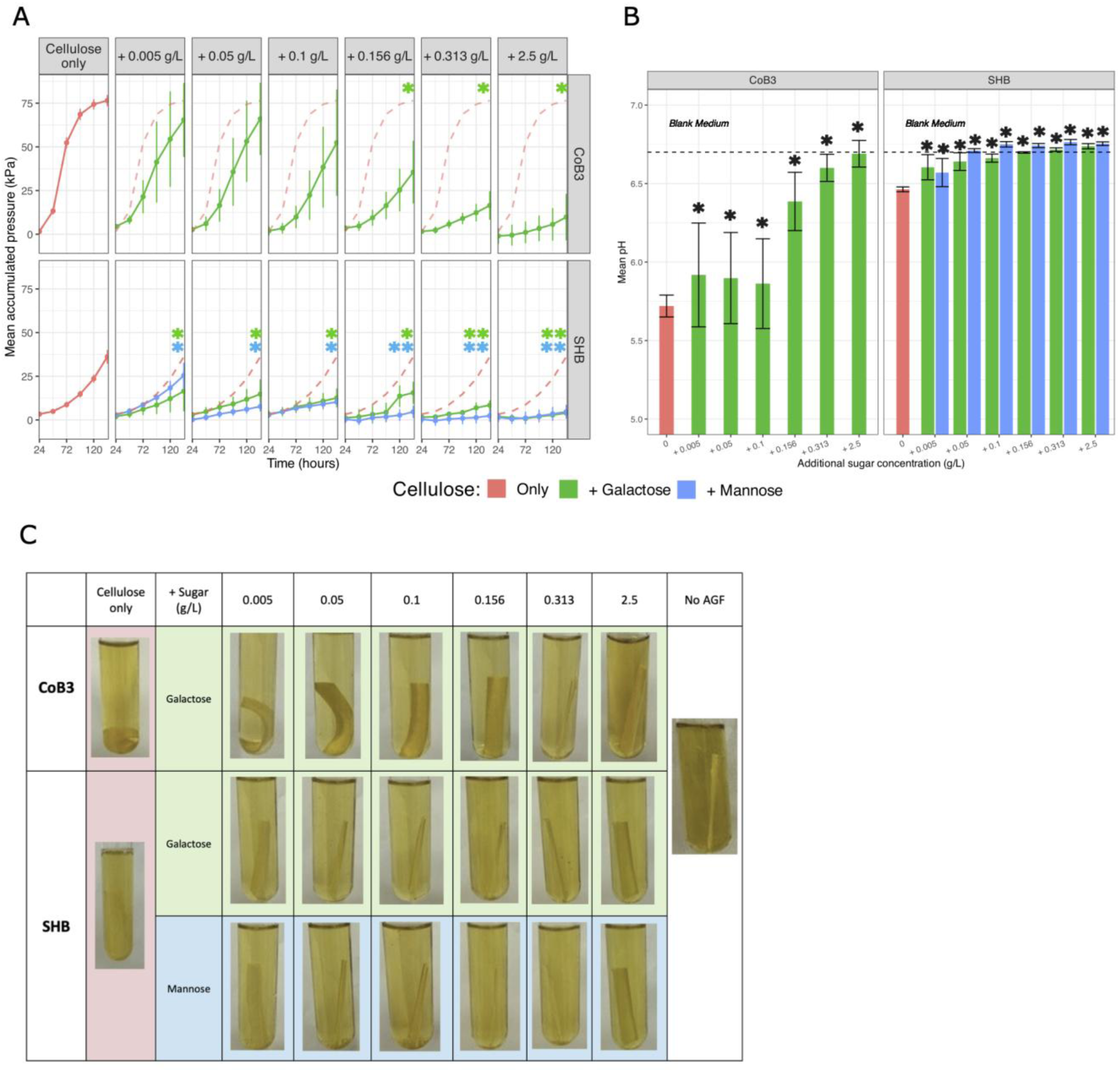
Sensitivity of growth of N. frontalis CoB3 and C. communis SHB on filter paper to inhibitory sugars at decreasing concentrations. Filter paper (1 % (w/v)) supplemented with galactose (CoB3 and SHB) or mannose (SHB) at a range of dilutions encompassing 2.5 g L^-1^ to 0.005 g L^-1^. Values reported are mean ± stdev (n=5). A) Fermentation gas accumulation of cultures and B) pH of culture supernatant after 144 h after fungal inoculation. C) Photographs of representative cultures at 144 h after fungal inoculation. Significance values reported are for AGF isolates on filter paper + additional sugar cultures versus cellulose-only cultures; one-way ANOVA and Tukey HSD analysis was used to assess CoB3 gas pressure and pH of CoB3, and for SHB gas pressure and both isolates’ pH Kruskal-Wallis and pairwise Wilcoxon test was used. For all statistical tests p-values are given: *** p < 0.001; ** 0.001 ≤ p < 0.01; * 0.01 ≤ p < 0.05, ns p ≥ 0.05.

*N. frontalis* CoB3 growth was statistically impeded by the addition of galactose at concentration of 0.156 g L^-1^ and higher, assessed by comparing total gas pressure accumulation and pH (Fig. 3). However, at lower concentrations (0.005 g L^-1^ – 0.1 g L^-1^), there was large variation amongst the biological replicates for both the total accumulated fermentation gas and pH of cultures measured, suggesting that the observed CoB3 growth at these concentrations of was unstable when compared to cultures in which filter paper was the sole carbon source (Fig. 3). Based on fermentation gas production, the concentration threshold of complete growth inhibition lies between 2.5 g L^-1^ and 0.313 g L^-1^. However, whilst fermentation end products may not be significantly affected at low galactose conditions, the distinct curling of the filter paper during degradation was only partially present at 0.005 g L^-1^ (Fig. 3C), suggesting the degradation of filter paper may still be perturbed even at low galactose concentrations.

Together, these results indicate that both *N. frontalis* CoB3 and *C. communis* SHB are highly sensitive to galactose or both galactose and mannose, respectively, when present at culture inoculation.

### Inhibitory non-metabolisable sugars also affect established AGF cultures

To evaluate whether the observed sugar-mediated inhibition of fungal isolate growth is specific to the initial inoculation and colonisation of the filter paper or instead reflects a broader disruption of the degradative processes required for growth on cellulose, we explored the effects of galactose and mannose addition to exponential growth of *N. frontalis* CoB3 and *C. communis* SHB.

After treatment with either 2.5 g L^-1^ galactose or mannose, a hygromycin positive control for inhibition and mannitol negative control, cultures were followed for 5 days to confirm hygromycin-treated cultures fermentation gas production had ceased (Fig. 4A). For both CoB3 and SHB, the addition of galactose during exponential growth did not significantly affect the total fermentation gas accumulation (Fig. 4A,B). However, the addition of mannose did significantly reduce the total fermentation gas accumulation of SHB (Fig. 4A,B). This could suggest that these hexose sugars do not share a common mode of action for the inhibition of SHB growth. Despite no differences in fermentation gas production, *N. frontalis* CoB3 culture medium pH was significantly higher when galactose was added compared to filter paper as the sole carbon source (Fig. 4C). This shift could reflect the respective ratios of acidic end products or reduction in the total amount of fermentation products, suggesting the presence of galactose could alter CoB3 metabolic routing. The assessment of the effect of sugar addition to SHB culture medium pH was constrained, as the pH recorded for filter paper as the sole carbon source showed limited reduction compared to the blank medium (Fig. 4C). However, visual inspection of the filter paper did show reduced swelling when galactose or mannose were added, similar to the hygromycin treated cultures (Fig. 4D). This suggests that the observed swelling of the filter paper is not solely from residual or secreted CAZyme activity but may be directly related to the success of SHB growth which galactose and mannose may hinder. The addition of mannitol, a sugar alcohol chosen for its similar structure to sugars allowing a comparable osmotic effect without introducing a metabolisable carbon source (Fig. S4), confirmed the results measured for both isolates were specific to the sugar treatments and not a result of osmotic stress (Fig. 4).

**Figure 4.**
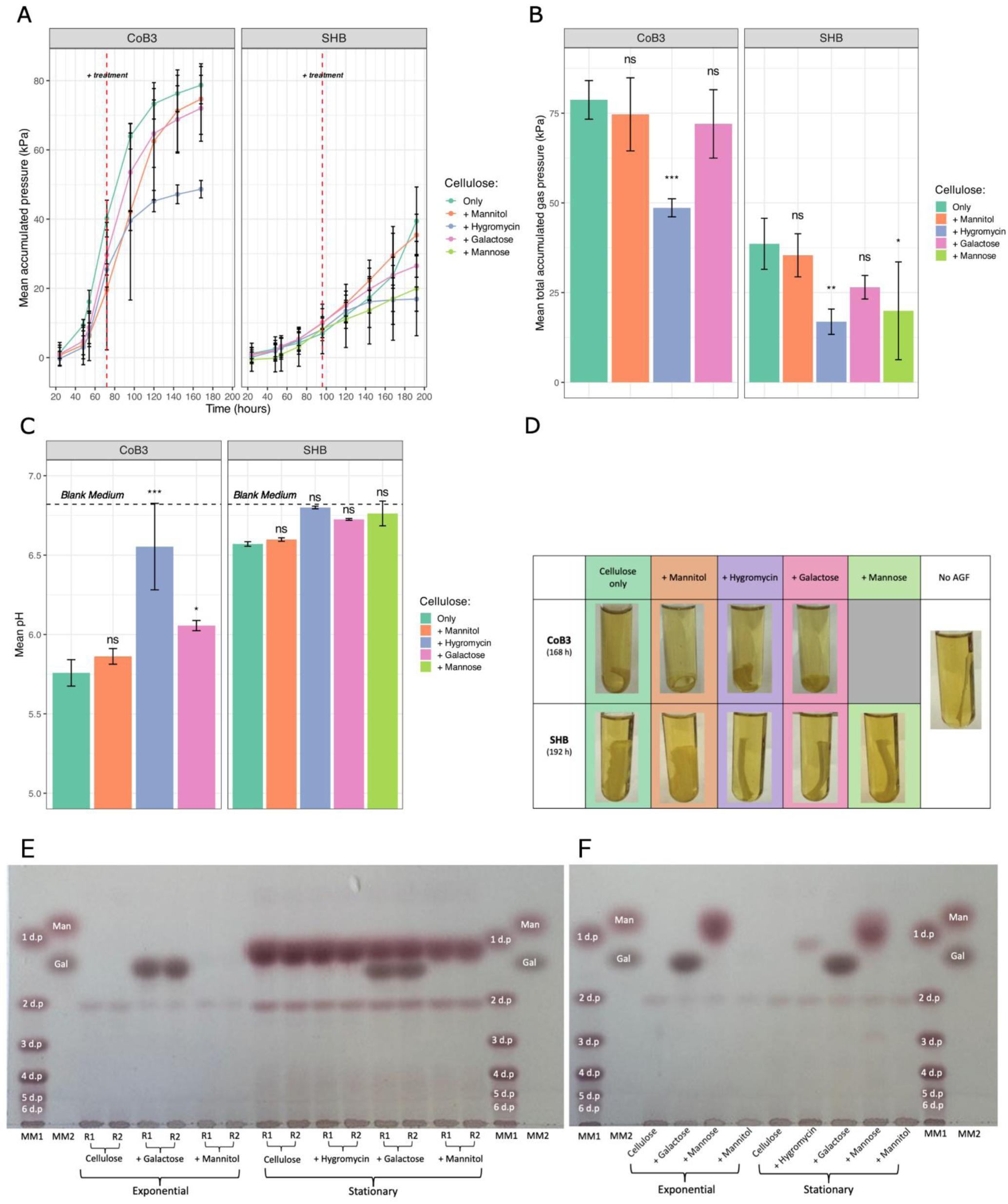
Growth of N. frontalis CoB3 and C. communis SHB on filter paper, with treatments added during exponential growth. Values reported are mean ± stdev (n=5). A) Fermentation gas accumulation of cultures, with dashed line indicating time point of treatment with galactose, mannose, mannitol (all to a final concentration of 2.5 g L^-1^) or hygromycin (to 150 µg mL^-1^), B) total accumulated fermentation gas, and C) pH at 168 h (CoB3) 192 h (SHB) after fungal inoculation. D) Photograph of representative culture after last gas pressure measurement. For each condition, a representative replicate of the culture medium from each AGF isolate is shown analysed by thin-layer chromatography, including a sub-sample of the exponential growth supplemented with galactose (CoB3 and SHB) and mannose (SHB) to a final concentration of 2.5 g L^-1^ of E) N. frontalis CoB3, F) C. communis SHB. Marker mixture 1 (MM1) was a cello-oligosaccharide ladder (DP 1 to 6) and marker mixture 2 (MM2) was galactose and mannose, dissolved in water. Significance values reported are for AGF isolates on filter paper + treatment cultures versus cellulose-only cultures; one-way ANOVA and Tukey HSD analysis was used to assess total gas pressure and pH of CoB3, and for SHB pH Kruskal-Wallis and pairwise Wilcoxon test was used. For all statistical tests p-values are given: *** p < 0.001; ** 0.001 ≤ p < 0.01; * 0.01 ≤ p < 0.05, ns p ≥ 0.05.

**Figure 5.**
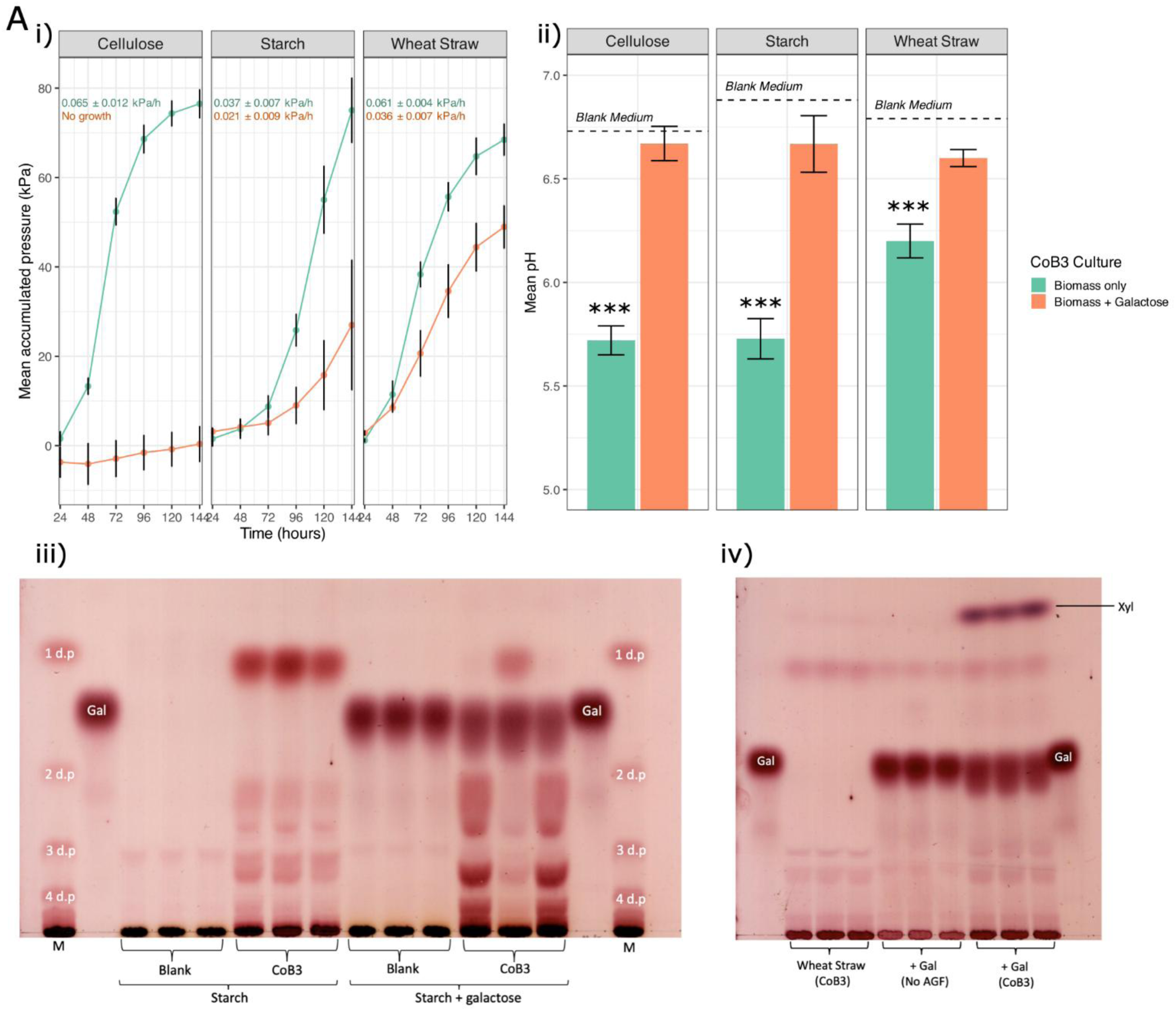
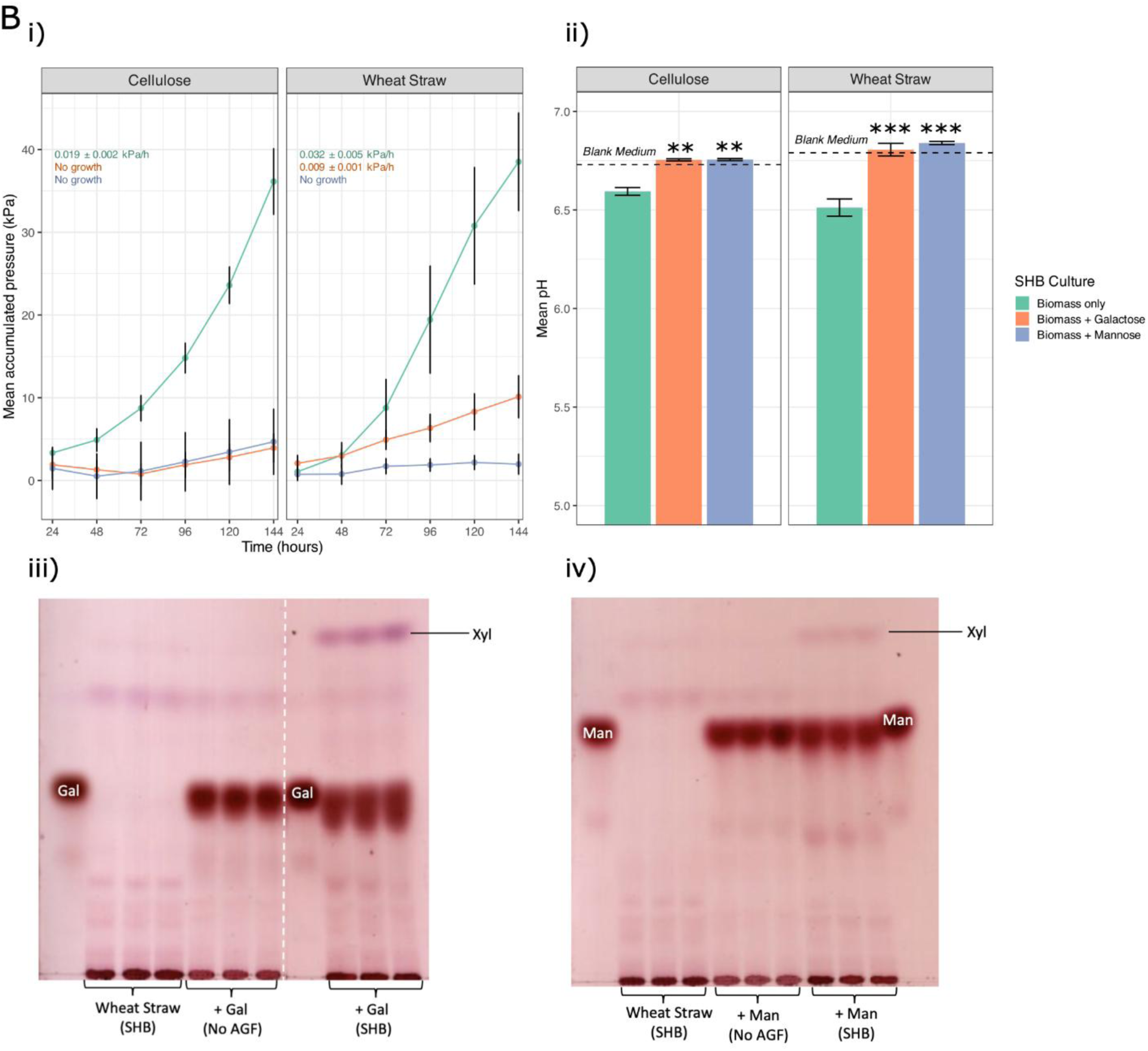
Growth of N. frontalis CoB3 and C. communis SHB on complex biomass carbon sources with addition of inhibitory sugars. A) N. frontalis CoB3 grown on filter paper, soluble potato starch or milled wheat straw. i) Fermentation gas accumulation of cultures (turquoise = biomass only; brown = biomass + 2.5 g L^-1^ galactose), and ii) pH at 144 h after fungal inoculation. TLC analysis of iii) soluble potato starch cultures and iv) milled wheat straw cultures. B) C. communis SHB grown on filter paper or milled wheat straw supplemented with galactose or mannose where indicated (2.5 g L⁻¹). i) Fermentation gas accumulation of cultures (turquoise = biomass only; brown = biomass + 2.5 g L^-1^ galactose; blue, biomass + 2.5 g L^-1^ mannose), and ii) pH after 144 h after fungal inoculation. TLC analysis iii) galactose-supplemented and iv) mannose-supplemented cultures. For both AGF isolates, marker lanes included galactose and mannose dissolved in water or a malto-oligosaccharide ladder (M; DP 1–6, 0.1% w/v), as appropriate. Significance values reported are for AGF isolates complex biomass + treatment versus complex biomass only cultures; Student’s t-test was used to assess total gas pressure and pH; p-values are given: *** p < 0.001; ** 0.001 ≤ p < 0.01; * 0.01 ≤ p < 0.05, ns p ≥ 0.05.

TLC analysis of the culture medium sampled did not show visible uptake of galactose by either AGF isolate, or mannose by *C. communis* SHB (Fig. 4E,F). Mannitol could not be visualised here, as the thymol-sulphuric acid stains sugars by the acidification of their active ketone/ aldehyde groups, which as a sugar alcohol mannitol does not possess (Jork *et al.,* 1990). At exponential growth, before the addition of the treatments, both AGF isolates produced a dimer degradation product (likely to be cellobiose) (Fig. 4E,F). This product was also present at a similar concentration in the culture medium at the end-point sampling of both isolates (Fig. 4E,F), with CoB3 also producing a glucose product (Fig. 4E). Comparison between CoB3 culture treatment groups of end point samples showed no differences in digestion products visualised by TLC, including the hygromycin control (Fig. 4E). As the treatments were added during exponential growth, it is possible that the CAZymes were already secreted into the culture medium which produced the dimer and glucose products, and their uptake could have been limited by depletion of another nutritional requirement from the culture medium or by the decrease in culture pH by volatile fatty acid (VFA) production. Whilst SHB also showed no effect on products observed in the end point samples for most treatments, the addition of hygromycin did show an accumulation of glucose (Fig. 4F), suggesting that galactose and mannose may have an inhibitory mechanism different from that of hygromycin.

Overall, whilst the addition of galactose did perturb visual degradation of filter paper for both *N. frontalis* CoB3 and *C. communis* SHB, their exponential growth is seemingly robust enough to produce similar quantities of fermentation end products.

### Effect of sugar addition on utilisation of alternative complex biomass carbon sources

Next, we investigated the effect of galactose and mannose supplementation to both fungal isolates’ growth on wheat straw, and for *N. frontalis* CoB3 on soluble potato starch (*C. communis* SHB cannot utilise starch to support its growth). This was to determine whether the inhibitory effects of soluble sugars observed at culture inoculation were specific to the colonisation and degradation of cellulose itself or instead reflected a conserved response during complex biomass colonisation.

The addition of galactose to cultures containing starch or wheat straw had an inhibitory effect on *N. frontalis* CoB3 growth, indicated by reduced accumulation of fermentation gases (*p*-value < 0.001) and increased culture medium pH compared with each respective complex biomass as the sole carbon source (Fig. 5A i, ii). However, complete growth inhibition was only observed when filter paper was the primary carbon source. TLC analysis of the culture medium revealed that galactose remained in the medium at a similar concentration when added to starch or wheat straw (Fig. 5A iii, iv). Low concentrations of wheat straw digestion products were detected, but there was clear accumulation of xylose in galactose-supplemented cultures (Fig. S5). *N. frontalis* cultures on starch produced oligosaccharides and glucose, and galactose supplementation in two of the three biological replicates resulted in higher oligosaccharide concentrations and a reduction in glucose, suggesting reduced growth in these replicated and delayed degradation of starch to glucose.

As observed for *N. frontalis* CoB3, galactose supplementation significantly reduced *C. communis* SHB growth on wheat straw, indicated by the reduced total accumulation of fermentation gas (*p*-value < 0.001) and increased culture medium pH compared with the non-supplemented culture (Fig. 5B i,ii). In addition, as observed for filter paper, mannose supplementation also resulted in complete growth inhibition when wheat straw was the primary growth substrate (Fig. 5Bi, ii). TLC analysis revealed, much like CoB3, accumulation of xylose when these sugars were present (Fig. 5B iii, iv). The xylose accumulation seemed proportional to the AGF growth, with CoB3 showing the highest xylose accumulation, followed by SHB when supplemented with galactose, with only a faint band of xylose observed for SHB when supplemented with mannose (Fig. 5).

These results reveal that the presence of galactose and/or mannose slows growth and perturbs the degradative activity of *N. frontalis* CoB3 and *C. communis* SHB on complex biomass substrates.

### Plant-derived polymers affect AGF growth on filter paper

As mono-and disaccharides affected fungal growth on the filter-paper model of wall cellulose, we also assessed the effects of supplementation with selected plant-derived polymers relevant to agricultural ruminant diets. This was to determine whether inhibitory effects associated with e.g. galactose and mannose are specific to their free sugar forms or are also exhibited when components of polysaccharides.

Whilst both fungal isolates failed to grow when supplemented with lignin extracts and pectin, possibly due to unforeseen interactions of these polymers with culture medium buffering capacity (Fig. 6A), both AGF isolates showed markedly different responses to the presence of other polymers, as observed with the addition of free sugars (Fig. 2). *C. communis* SHB growth as measure of fermentation gas accumulation was perturbed by the addition of the polymers, whereas *N. frontalis* was unaffected as measured by fermentation gas production and culture medium pH (Fig. 6A). The inhibitory effects of galactose and mannose on *C. communis* SHB growth was conserved in their polymeric forms, as the addition of both galactomannan and glucomannan resulted in completely suppressed SHB growth (Fig. 6A). However, *N. frontalis* CoB3 was not significantly affected by the presence of galactomannan, suggesting the inhibitory effect of galactose on this isolate is contingent on the physical state of the sugar (free versus polymeric), as the presence of mannose was confirmed not to be facilitating growth recovery (Fig. S6). Unexpectantly, the addition of arabinoxylan had a beneficial effect on SHB growth, significantly increasing fermentation gas accumulation (*p-*value < 0.001) and decreasing culture pH.

**Figure 6.**
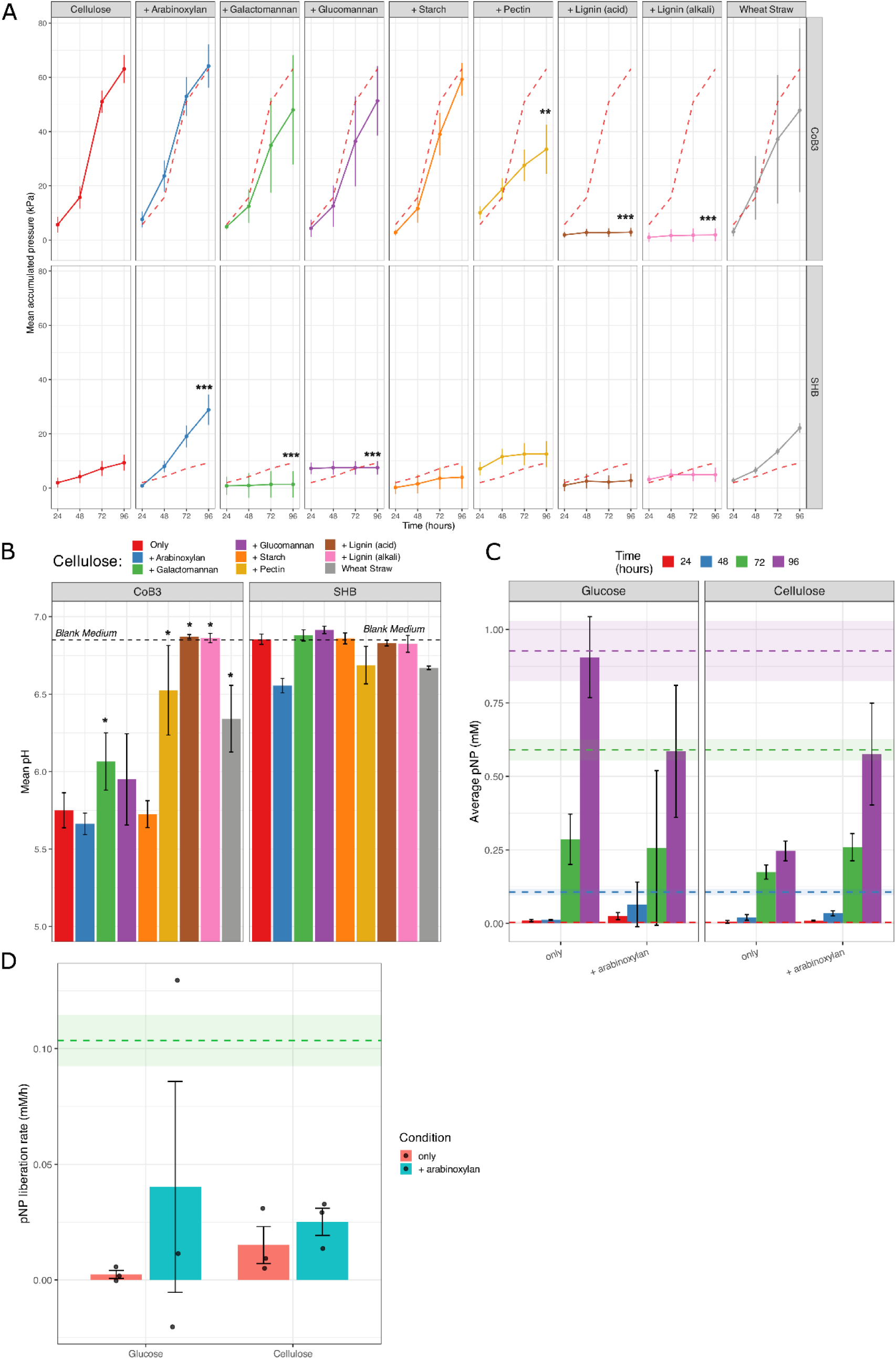
Effect of addition of individual polymers on fungal growth and enzyme activity. Filter paper supplemented with isolated distinct polymers. A) Fermentation gas accumulation of cultures, B) pH of culture at 144 h after fungal inoculation; the dashed line represents the pH of blank culture medium. Values reported are mean ± stdev (n=5). Significance values reported are for AGF isolates on filter paper + supplementary polysaccharide cultures versus cellulose-only cultures; Kruskal-Wallis and pairwise Wilcoxon test was used to assess CoB3 gas pressure and both isolates pH, and a one-way ANOVA and Tukey HSD analysis was used to assess SHB gas pressure. N. frontalis CoB3 cultures were assessed for xylosidase activity in response arabinoxylan supplementation when grown on glucose or filter paper by; C) pNP (mM) liberated from xylo-pNP every 24 h and D) the average rate of pNP liberation from xylo-pNP over the 96 h growth period for each culture. The wheat straw control average and standard deviation shown by dashed lines and coloured stripe respectively. Values reported are mean ± stdev (n=3). Significance was assessed using a two-way ANOVA and Tukey HSD analysis.

As AGF may regulate CAZyme expression independently of their overall fermentation output (Brown *et al.,* 2023), *N. frontalis* CoB3 may be responding to the presence of additional polymers by inducing degradative enzymes despite no significant difference being measured in its fermentation gas production or culture medium pH (Fig. 6A). Therefore, as *Neocallimastix* species have been reported to encode an abundance of xylanases in their genomes (Solomon *et al.,* 2016), *N. frontalis* CoB3 xylosidase activity was examined to determine whether the presence of arabinoxylan elicited an inducible degradative response.

Overall, the addition of arabinoxylan was shown to increase *N. frontalis* CoB3 xylosidase activity compared to each respective sole carbon source (Fig. 6D). The increase of xylosidase activity in response to the added arabinoxylan was shown to be more tightly conserved across biological replicates when filter paper was the primary substrate compared to when it was glucose, and cellulose elicited clearer trends consistent with substrate-specific induction (Fig. 6C). The widely variable xylosidase activity across biological replicates grown on glucose is unlikely to be directly correlated with the rate of glucose consumption, as this was the same across replicates regardless of arabinoxylan supplementation (Fig. S7). The variable measured xylosidase activity for cultures grown on glucose could be a starvation response and a scavenging strategy in which there is a basal xylosidase expression regardless of growth substrate (Gomez, Durand and Fèvre, 1998), to prime the fungal cells for the colonisation of plant biomass (Butkovich *et al.,* 2025; van Munster *et al.,* 2014). Therefore, filter paper can be used as a physiologically relevant model for interrogating substrate-induced enzyme responses, especially for polysaccharides that do not provide metabolisable sugars to AGF isolates.

## Discussion

Whatman No.1 filter paper is composed almost entirely of cellulose and is largely free of cutin and hemicelluloses. It also possesses a highly crystallinity index of approximately 80% (Costa *et al.,* 2014; Ferrara *et al,* 2024; Sandy, Manning and Bollet, 2010). Consequently, it has frequently been used as a cellulose substrate for assessing microbial cellulase activity and growth (Bauchop, 1981; Guder and Krishna, 2019; Lowe, Theodorou and Trinci, 1987; Sijpesteijn, 1975). Here, *N. frontalis* CoB3 and *C. communis* SHB could both grow successfully on filter paper relative to their respective growth on lignocellulosic material (Fig. 1). Furthermore, these isolates exhibited vastly different physical effects on the cellulose: *N. frontalis* CoB3 growth caused the filter paper to curl whereas *C. communis* SHB growth swelled it. *N. frontalis* CoB3 is of the more commonly reported rhizoidal morphology, which allows it to penetrate and anchor into plant material whereas *C. communis* SHB has the less common bulbous morphology, forming spherical sporangia (Shen *et al.,* 2026). Previous characterisation of another bulbous isolate, *Caecomyces communis var. churrovis* has led to hypothesis that bulbous AGF isolates compensate for their lack of rhizoids by secreting free CAZymes to break down polymeric substrates (Henske et al., 2017). Therefore, the differences observed on the filter paper may be linked to the differing physical and enzymatic capabilities driven by the two cultures’ differing morphologies.

The addition of free sugars revealed further differences in fungal isolate responses when filter paper was the primary carbon source. The presence of glucose and cellobiose both increased *C. communis* SHB growth rate (Fig. 2), contrary to the expected effect of the presence of metabolisable free sugars inducing carbon catabolite repression and reducing filter paper degradation (Adnan *et al.,* 2017). Therefore, it is possible the rate of cellulose digestion limits the growth of this isolate. Supplementation with galactose was inhibitory to both isolates’ growth, and mannose was also inhibitory to the growth of SHB (Fig. 2). This was unexpected, as previous screening of these isolates’ responses to galactose had shown no measurable response when glucose was the primary carbon source (Matthews *et al.,* 2026). In addition, the synergistic co-substrate utilisation of glucose and mannose by *N. frontalis* CoB3 (Matthews *et al.,* 2026) producing increased fermentation gas yields was not observed here. These findings suggest that the metabolic accessibility of the primary substrate influences the fungal responses to the presence of additional sugars, but the mechanism behind this remains unclear. As the sugars shown here to repress *N. frontalis* CoB3 and *C. communis* SHB growth are epimers of glucose, it may be possible that these sugars can disrupt key metabolic pathways required for successful growth on filter paper, owing to their structural similarity to glucose. Comparable effects have been reported for rare sugars, where structurally related monosaccharides interfere with established metabolic pathways, for example D-arabinose and D-tagatose have been shown to inhibit growth or physiological processes in the roundworm *Caenorhabditis elegans* (Sakoguchi *et al.,* 2016) and in the bacterium *Streptococcus mutans* (Hasibul *et al.,* 2017), suggesting that analogues of common sugars can perturb sugar metabolism even when they are not efficiently utilised as carbon sources.

The inhibitory effect of galactose when filter paper is the primary carbon source seems to be a conserved response over different AGF genera and families, as this was previously observed in a similar screening (Henske *et al.,* 2018). However, notably, the concentration of supplemented sugars was four times higher, and the media were inoculated with only zoospores (Henske *et al.,* 2018a), whereas cultures here were inoculated by a routine sub-culture. Sugar sensing and signalling pathways are often integrated with fungal cellular physiological states to enable the regulation of their growth and development (Johns *et al.,* 2021; Rutherford *et al.,* 2019). Therefore, whilst it remains uncertain whether there are differing threshold-dependent responses to supplemented sugars between growth stages in AGF, a threshold-dependent response could potentially account for the biological variation of fermentation gas accumulation attributed to unstable growth recorded for *N. frontalis* CoB3 at the lower concentrations of supplemented galactose tested (Fig. 3). In addition, this could explain why complete growth inhibition was only observed when galactose or mannose were supplemented at fungal inoculation, and not when added during exponential growth (Fig. 4).

Alternatively, the inhibition of fungal growth by galactose may reflect toxic accumulation of galactose 1-phosphate (Gal-1-*P*), a known phenomenon in other fungi including *Aspergillus nidulans* and *Candida albicans* (Boulanger *et al.,* 2021), which can grow on galactose. In these systems, incomplete or imbalanced flux through the Leloir pathway (Gal → Gal-1-*P* ↔ UDP-Gal ↔ UDP-Glc; Sharples & Fry, 2007) results in Gal-1-*P* accumulation, leading to growth inhibition even when galactose is not utilised as a primary carbon source. From the sequenced genome of *N. frontalis* CoB3, this complete galactose catabolism pathway is present (https://mycocosm.jgi.doe.gov/Neofron1/Neofron1.home.html), despite the organism not being able to utilise galactose as the sole carbon source (Matthews *et al.,* 2026). Furthermore, the Leloir pathway has previously been identified at the transcriptomic level in a diverse range of AGF isolates, including those belonging to the genera of *Neocallimastix* and *Caecomyces* (Murphy *et al.,* 2019), so it is likely *C. communis* SHB also possesses this pathway despite it also being unable to utilise galactose for growth (Matthews *et al.,* 2026). It is thought that the Leloir pathway identified in these AGF was acquired via horizontal gene transfer (Murphy *et al.,* 2019). Therefore, this pathway may be missing the required promoters or transcription factors to be active and properly regulated in AGF. As a result, the inhibitory effect of galactose when present at fungal inoculation could be due to an accumulation of intracellular Gal-1-*P* that cannot be effectively detoxified, whereas during exponential growth, the established metabolic activity may mitigate this toxicity through dilution effects or an ability to export metabolic intermediates. This could explain the lack of growth inhibition when galactose was added during exponential growth and the observed change in pH of CoB3 when galactose was added at exponential growth compared with filter paper as the sole carbon source (Fig. 4C). Whilst TLC of culture medium of both isolates did not reveal detectable galactose uptake (Fig. 4E,F), the sensitivity of this approach may be insufficient to capture low-flux or transient uptake events, as Gal-1-*P*-mediated toxicity in *S. cerevisiae* mutants could be achieved at the equivalent 0.02 g L^-1^ (Mumma *et al.,* 2008).

It is uncertain whether *C. communis* SHB has a complete mannose catabolic pathway; however, a similar accumulation of mannose 6-phosphate (Man-6-*P*) could result in a similar metabolic stress, inhibiting growth (Boulanger *et al.,* 2021). Previously, *N. frontalis* CoB3 was shown to be able to metabolise mannose in combination with glucose (Matthews *et al.,* 2026), therefore, may be able to cope with the generation of Man-6-*P* when utilising cellulose for growth.

Although galactose was inhibitory to AGF growth on alternative complex biomass (straw or starch), it did not completely repress fungal growth as observed on filter paper (Fig. 5). The mitigation of complete repression observed by *N. frontalis* CoB3 during the utilisation of starch is likely to be a result of this isolate employing distinct degradative mechanisms, signalling pathways, and metabolic fluxes compared to those involved in filter paper utilisation. As ability to utilise starch as the sole carbon source is not conserved across the Neocallimastigomycota phylum (Gordon and Phillips 1998), it is unclear how representative this observed response is. Likewise, both isolates’ growth on wheat straw was hindered but not completely repressed by supplementation with galactose (Fig. 5). As lignocellulose is the preferred native substrate of AGF (Akin and Borneman, 1990), it is possible that the presence of additional polymers in wheat straw may trigger alternative metabolic pathways which can compensate for the inhibitory effects of galactose, resulting in sub-optimal growth (Fig. 5).

Mannose suppressed *C. communis* SHB growth even when cultures were grown on wheat straw (Fig. 5), indicating that this response persists on complex lignocellulosic substrates. The use of filter paper as a simplified model of the lignocellulosic backbone enabled the investigation of fungal growth responses to isolated carbohydrates of interest, including those which cannot independently support AGF isolates’ growth. In this model, the inhibitory effects of mannose (and galactose) were retained when supplied in their polymeric forms, as supplementation of galactomannan and glucomannan both resulted in complete suppression of *C. communis* SHB growth (Fig. 6A). Consistent with these observations, mannan-containing polysaccharides have been shown inhibit cellulase production in several filamentous fungi, including *Neurospora crassa, Trichoderma reesei*, and *Aspergillus* spp. (Hassan *et al.,* 2019). This inhibition has been attributed to crosstalk between mannan-and cellulose-responsive pathways, whereby manno-oligosaccharides and mannose interfere with cellulose signalling and disrupt cellulase induction, which could link to the impaired cellulose utilisation and inhibited *C. communis* SHB growth noted here (Fig. 6A). In contrast, *N. frontalis* CoB3 did not exhibit inhibition by galactomannan (Fig. 6A), which may reflect differences in substrate sensing, regulatory crosstalk, or metabolic capacity between the isolates. Whilst CoB3 did not show any responses to the additional polymers by the parameters of fermentation gas accumulation and culture medium pH (Fig. 6A), it was shown to respond to the presence of arabinoxylan by increased production of xylosidases (Fig. 6B).

Overall, these findings demonstrate species-specific differences in AGF degradation strategies, highlighting the need for better characterisation of AGF metabolic capabilities and CAZyme repertoires for their application in lignocellulosic bioconversion and ruminant nutrition. The identification of galactose and mannose as inhibitory to AGF isolates is particularly relevant in relation to rumen microbiome modulation strategies, where dietary supplements for the early-life programming of the rumen include galactose-and mannose-oligosaccharides fed either directly to the calves by being dissolved in milk replacers or as a top-dressing on solid feed starter, or indirectly through maternal supplementation if the young is fed directly from the mother (Chang et al., 2022) (Hu et al., 2024). Such interventions could disrupt AGF establishment and reduce fibre degradation efficiency, a key parameter in achieving efficient production of these animals.

## CRediT authorship contribution statement

**JLM:** Methodology, Conceptualization, Investigation, Data curation Visualization, Validation, Formal analysis, Funding acquisition, Writing – original draft, Writing – review & editing. **SCF:** Conceptualization, Supervision, Project administration, Funding acquisition, Writing – review & editing. **JMvM:** Conceptualization, Methodology, Formal analysis, Data curation, Supervision, Project administration, Funding acquisition, Writing – review & editing.

## Acknowledgements

The authors are grateful for support from the Royal Society via URF\R1\231686, and from the UKRI Biotechnology and Biological Sciences Research Council (BBSRC) through the EASTBIO DTP, grant number BB/T00875X/1.

