## Supplementary Material for "Sugar-mediated inhibition of growth and lignocellulose degradation in anaerobic gut fungi revealed using cellulose filter paper"

### Supplementary information

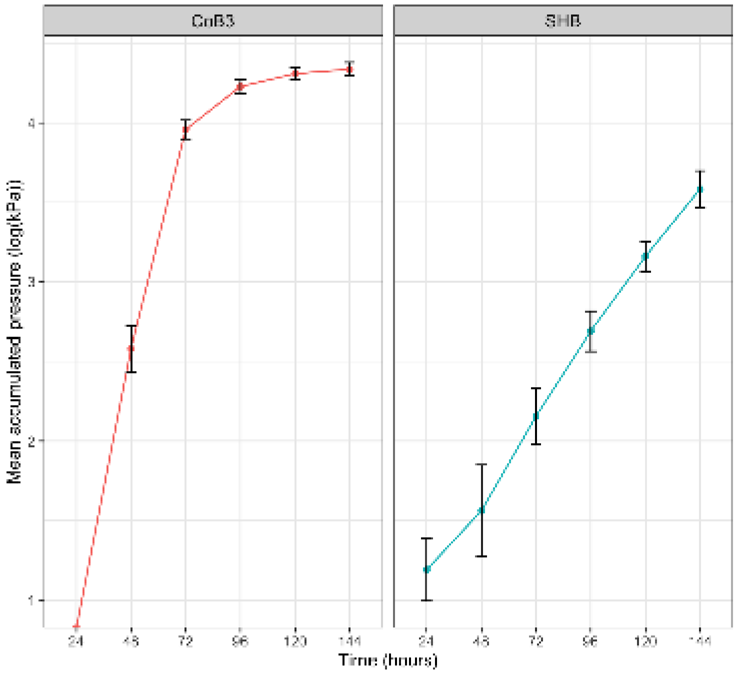


***Supplementary Figure 1. Growth curve of log*** $_{10}$ ***transformation of mean accumulated gas pressure of AGF isolates used to identify exponential growth stage on filter paper.***

Exponential growth rates were identified from the $\log_{10}$ transformed mean accumulated gas pressure of AGF isolates and visual identification of the region of linearity. To this region, linear regression was then applied, and the resulting slope was used to estimate the exponential growth rate. Here, for example, the region of linearity was identified for *N. frontalis* CoB3 to be 24 to 72 hours and for *C. communis* SHB 48 to 144 hours. Linear regression was applied to these respective regions, the calculate an estimated average exponential growth rate of 00065 ± 0.008 kPa/h for *N. frontalis* CoB3 and 0.0021 ± 0.003 kPa/h for *C. communis* SHB.

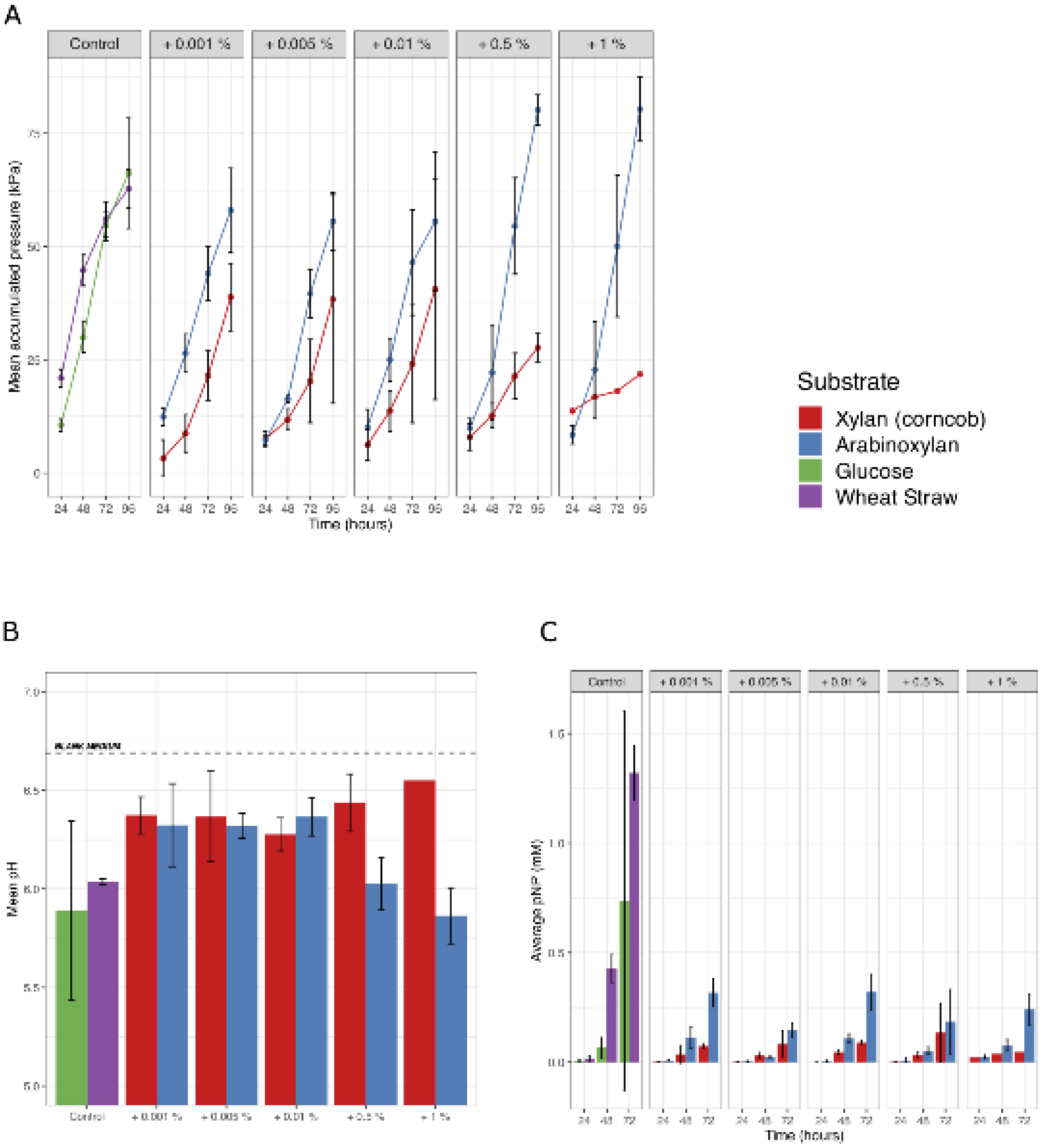


***Supplementary Figure 2. Trial of additional polysaccharide concentration using N. frontalis CoB3***

*Glucose (5 g $L^{-1}$) was supplemented with xylan from corncob or wheat arabinoxylan (1% to 0.001 %, w/v), and milled wheat straw (0.5 mm) (1 %, w/v). Values reported are mean ± stdev (n=5). A) Fermentation gas accumulation of cultures, and B) pH of culture supernatant after 96 h after fungal inoculation. C) Average pNP (mM) liberated from xylo-pNP every 24 hours.*

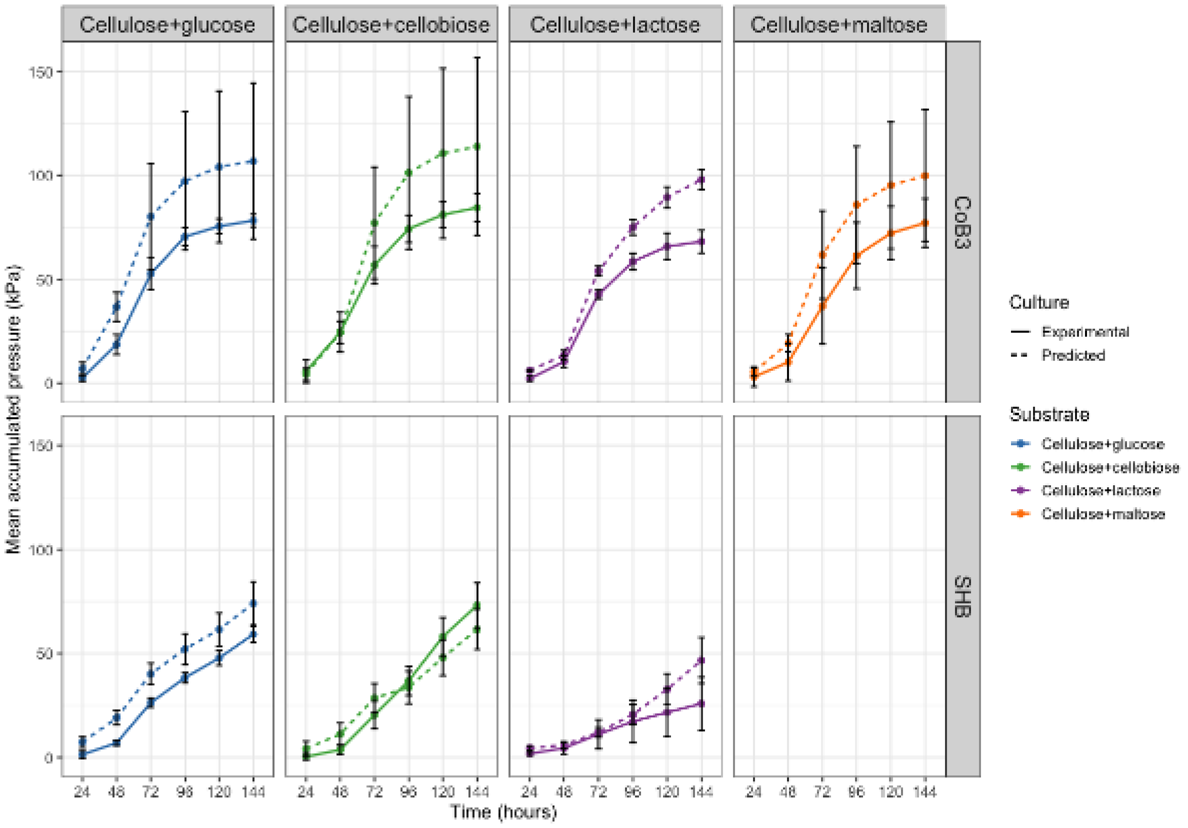


***Supplementary Figure 3. Predicted growth curve of fungal isolates N. frontalis CoB3 and C. communis SHB on filter paper + metabolisable sugar.*** *As C. communis SHB cannot grow on maltose, this was not calculated.*

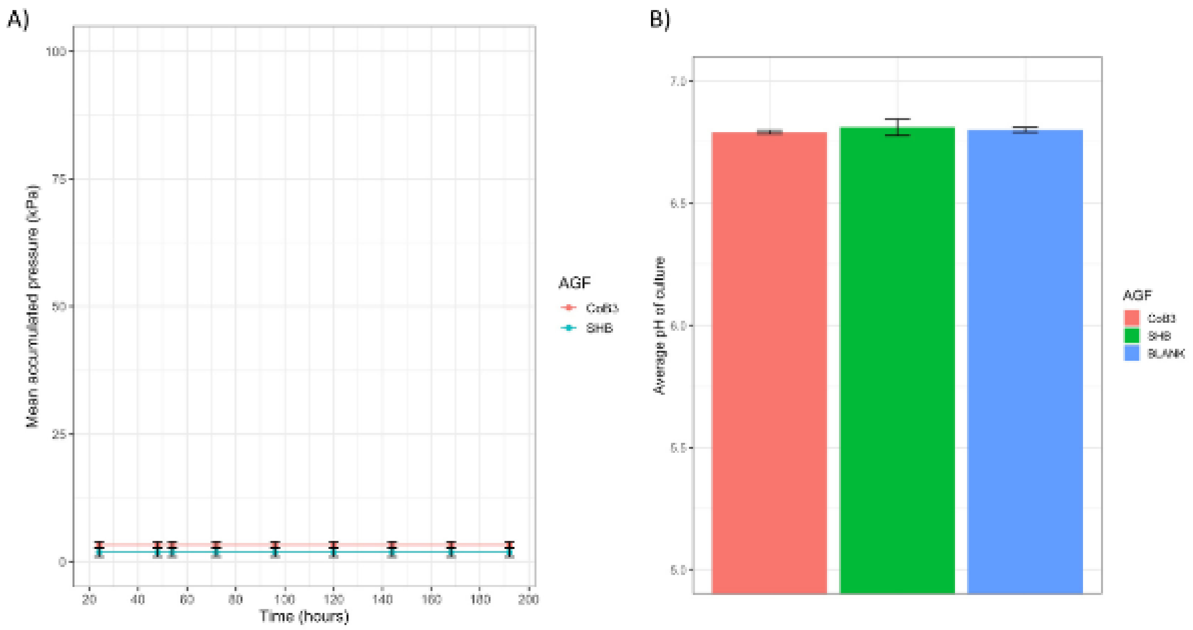


***Supplementary Figure 4. Growth curve of fungal isolates N. frontalis CoB3 and C. communis SHB on mannitol.*** *A) Fermentation gas accumulation by cultures grown on mannitol (5 g L$^{-1}$) and B) pH of culture supernatant after 144 h after fungal inoculation, 'BLANK' was medium without fungal inoculation.*

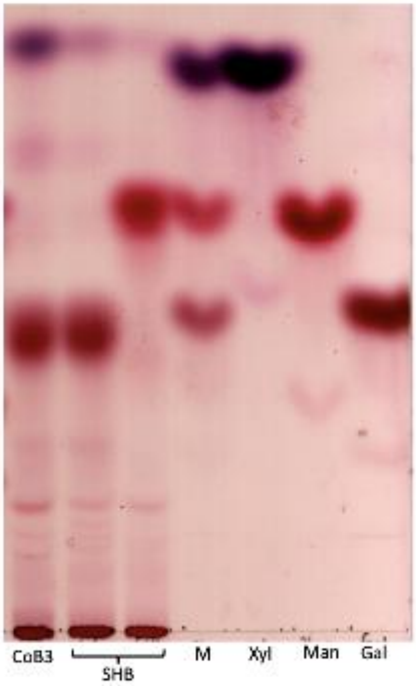


***Supplementary Figure 5. Confirmation by thin-layer chromatography of the liberation of xylose from wheat straw when supplemented with additional sugars.*** *For each wheat straw and additional soluble sugar condition, a representative replicate of the culture medium from each AGF isolate is shown analysed by TLC. Marker sugars; galactose, mannose, and xylose were loaded as a mixture and separately, dissolved in water.*

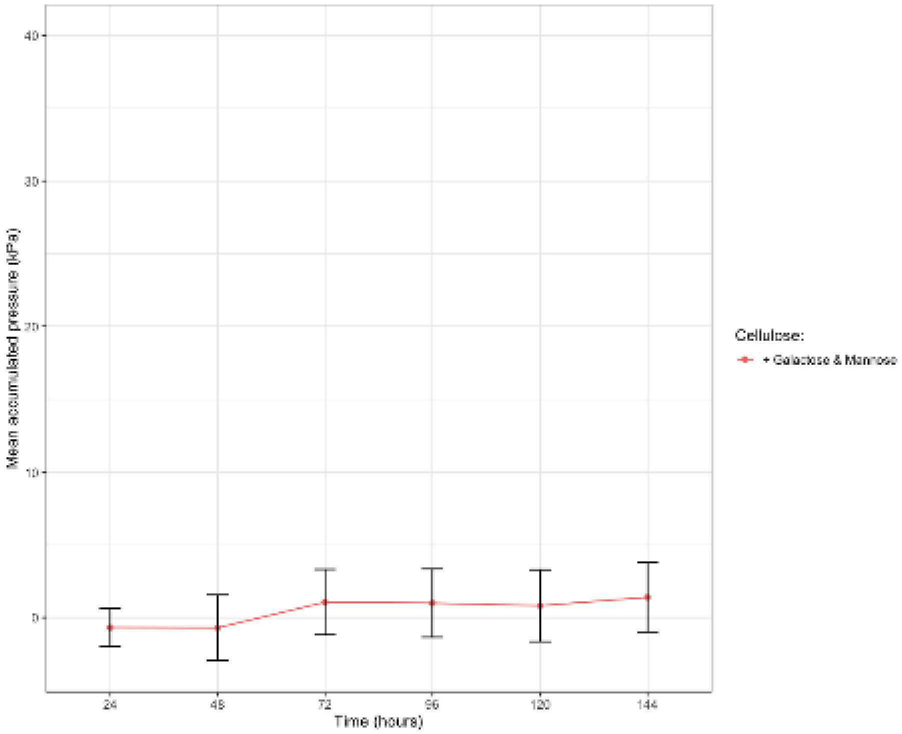


***Supplementary Figure 6. Growth curve of N. frontalis CoB3 on filter paper (1 %, w/v) and galactose and mannose.*** *To assess potential recovery effect on N. frontalis growth by the presence of mannose, both galactose and mannose added to the same ratios present when guar galactomannan was supplemented at 0.5% (w/v) (Table 1).*

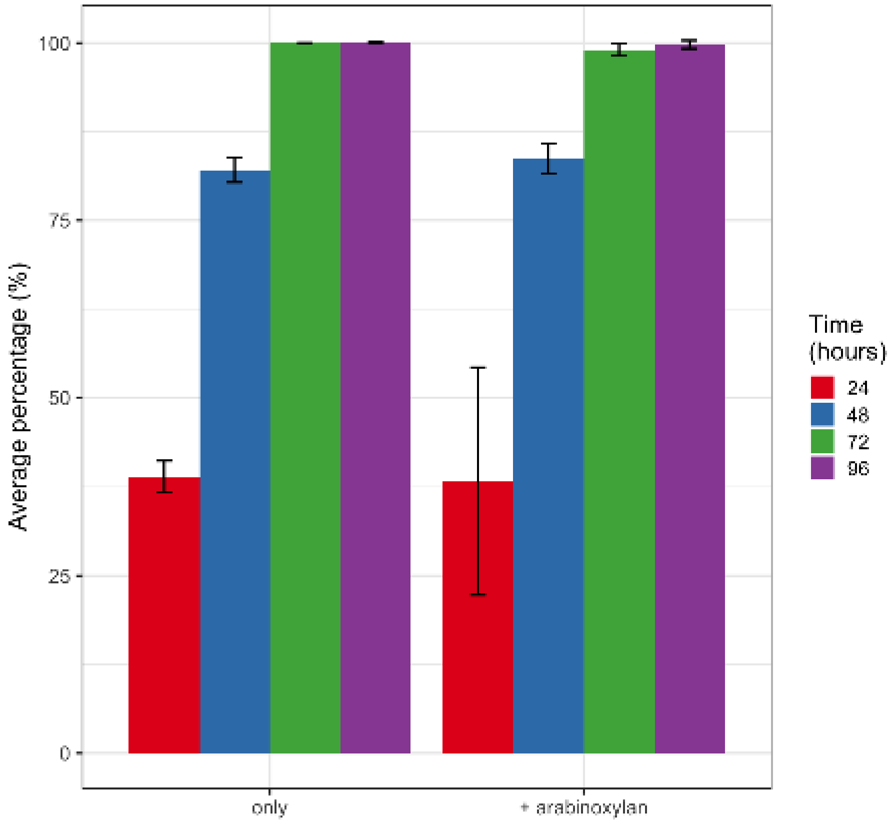


***Supplementary Figure 7. Average percentage of glucose removed from N. frontalis culture medium.*** *The baseline substrate was glucose (5 g L$^{-1}$) (only) and supplemented with wheat arabinoxylan (0.5 %, w/v). Values reported are mean ± stdev(n=5).*

Every 24 hours after gas pressure was recorded, 0.1 mL of culture medium was removed and placed on ice. The concentration of glucose was measured using D-Glucose Assay Kit (GOPOD Format, Megazyme) following the manufacturer's instructions. Blank medium (no glucose or fungal inoculation) was used to correct for the culture medium colour.
